# RNA sequencing variants are enriched for eQTL in cattle tissues

**DOI:** 10.1101/2024.04.29.591607

**Authors:** Alexander S. Leonard, Xena M. Mapel, Hubert Pausch

## Abstract

Association testing between molecular phenotypes and genomic variants can help to understand how genotype affects phenotype. RNA sequencing provides access to molecular phenotypes such as gene expression and alternative splicing while DNA sequencing or microarray genotyping are the prevailing options to obtain genomic variants. Here we genotype variants for 74 male Braunvieh cattle from both DNA and deep total RNA sequencing from three tissues. We show that RNA sequencing calls approximately 40% of variants (7-10 million) called from DNA sequencing, with over 80% precision, rising to over 92% of variants called with nearly 98% precision in highly expressed coding regions. Allele-specific expression and putative post-transcriptional modifications negatively impact variant genotyping accuracy from RNA sequencing and contribute to RNA-DNA differences. Variants called from RNA sequencing detect roughly 75% of eGenes identified using variants called from DNA sequencing, demonstrating a nearly 2-fold enrichment of eQTL variants. We observe a moderate-to-strong correlation in nominal association p-values (Spearman ρ^2^∼0.6), although only 9% of eGenes have the same top associated variant. We also find several highly significant RNA variant-only eQTL, demonstrating that caution must be exercised beyond filtering for variant quality or imputation accuracy when analysing or imputing variants called from RNA sequencing.

## Introduction

High-throughput RNA sequencing (RNA-seq) has been frequently applied for measuring gene expression levels (1), assembling *de novo* transcriptomes (2), detecting copy number alterations (3), and identifying genomic variants that influence gene expression (4). Genotypes called from RNA-seq have also been used to determine population structure (5, 6). Historically, RNA-seq has been viewed as unreliable input for performing genetic variation calling for several reasons: RNA-seq typically targets fewer fragments than DNA sequencing, messenger RNA (mRNA) only produces transcripts in coding regions (∼1% of a mammalian genome), and a comparative lack of algorithmic development relative to DNA variant callers. Typical whole-genome sequencing (WGS)-based studies might use between 200-500 million reads (corresponding to approximately 10-fold to 25-fold coverage of a mammalian genome), while RNA studies aim for between 30 million reads to measure gene expression and 100 million reads to quantify alternative splicing events and map splicing QTL (sQTL) (7). While mRNA is the most widely used source for RNA-seq, total RNA, with pre-transcription RNA, contains more than just coding sequences including intronic and even intergenic regions (8). As a consequence of these factors, as well as expression variability in different tissues, far fewer genetic variants are called with RNA-seq than DNA-seq, with earlier studies identifying only 100k variants from 7 cow transcriptomes (9) or 68k variants from 29 cow transcriptomes (5). Even large studies like the cattle Genotype-Tissue Expression (GTEx) project (8), could only confidently call 22k variants from 7,180 publicly available transcriptomes of diverse origin, which is several orders of magnitude less than called from similar sized cohorts with WGS data (10). These variants can then be imputed to higher density using large reference panels, like that of the 1000 Bull Genomes project (10). However, a strong depletion of non-coding variants in typical RNA-seq datasets results in a less reliable imputation of variants that are distant to transcribed regions. Similar observations have been made in chicken (11), pig (12), and human (13).

RNA-seq variant callers are less common than their DNA-seq counterparts, but GATK (14) and a combination of preprocessing RNA-seq reads with Opposum (15) and calling variants with Platypus (16) have been the dominant options. Recently, DeepVariant has been extended to provide an RNA-seq trained model (17), greatly improving the accuracy and quantity of variants called from RNA-seq compared to the previous state of the art. The improved DeepVariant model also reduces the number of variants called at sites subjected to A-to-I editing within the RNA-seq. Such RNA editing events warrant attention as they can have important functional effects (18).

The mapping of expression and splicing quantitative trait loci (e/sQTL) is increasingly performed to investigate the impact of regulatory regions on phenotypes. These loci can be detected through association testing between molecular phenotypes (e.g., gene expression and splicing levels quantified from RNA-seq) and genetic variation. Recent studies have identified e/sQTL in cattle affecting economically relevant traits, such as male fertility (4), milk production (19), and carcass yield (20). These e/sQTL have proven highly valuable in prioritizing candidate causative variants for complex traits and diseases (8) (21).

In this work, we apply DeepVariant to call variants from deeper-than-usual total RNA sequencing. These RNA-seq based variant calls are enriched for eQTL and are strongly correlated for nominal p-values with the WGS-derived eQTL. We observe a substantial number of single nucleotide differences in DNA-RNA pairs and degradation of variant calling quality as total RNA-seq coverage drops to 100 million and 30 million reads.

## Methods

### DNA and RNA alignment

Publicly available DNA and RNA sequencing reads for three tissues for 74 samples from Mapel et al. (4) (Supplementary Table 3) were filtered with fastp (v0.23.4) (22). The DNA-seq data were aligned to the cattle reference (ARS-UCD1.2) with bwa-mem2 (v2.2.1) (23, 24) with the flag “-M”. The alignments were deduplicated and sorted with SAMtools (v1.19.1) (25). RNA-seq reads were aligned to the same reference and the Ensembl gene annotation (v108) using STAR (v2.7.9) (26) with --waspOutputMode and heterozygous SNPs from DNA to account for allelic imbalance. Read depth was estimated with perbase (v.0.8.5) (https://github.com/sstadick/perbase) and coverage per annotation classification was calculated with bedtools (v2.30) annotate (27) using the Ensembl v108 annotation.

Lower sequencing coverage was simulated by downsampling with SAMtools view -s <fraction>, where the fraction was chosen to approximately sample one hundred, thirty, and five million paired-end read subsamples.

### Variant calling and analysis

Variants were called from the aligned bam files using DeepVariant (v1.5) (28). For the DNA samples, we additionally used the “insert_size” channel, while for the RNA samples we used “--split_skip_reads” and the v1.4 RNA checkpoint model. All samples for each set of DNA or RNA tissue were merged using GLnexus (v1.4.1) (29). Variants were then phased and missing genotypes imputed using Beagle (v4.1) (30) using the “gl” field. Variant call intersection sets were calculated with BCFtools (v1.19) (36) isec, and precision/recall/F1 were calculated with hap.py (v0.3.15) (https://github.com/Illumina/hap.py), stratifying by region with a bed file containing annotated exon coordinates based on their expression level quantified in transcripts per million (TPM).

Principal components (PCs) were calculated with plink2 (v2.00a4LM) (31), using a minimum allele frequency of 5% and treating half calls as missing. Each individual’s breed was assigned according to the Swiss Braunvieh herdbook. Variant effects were classified with VEP (v108) (32), using the flags ‘--tab --fields “Consequence,IMPACT” --species bos_taurus’. Regions without variants were identified by convert VCF to BED format, followed by merging blocks within 1 Kb using bedtools merge. We then assessed uncovered regions using bedtools genomecov.

Allele-specific expression was calculated on the WASP filtered alignments with QTLtools (v1.3.1) (33) with the ase command, additionally including the flag “--both-alleles-seen” to remove monoallelic expression.

### eQTL analysis

We constructed TPM matrices (with filtering for WASP tag) using QTLtools quan and featureCounts (v2.0.4) (34), including genes with ≥0.1 TPM in ≥20% of samples and ≥6 reads in ≥ 20% of samples for association mapping. We then quantile normalised the TPM matrices. Principal components for LD-pruned variant calls and RNA expression were calculated with QTLtools pca.

We split multiallelic variants into multiple biallelic variants, followed by filtering variants at 1% minor allele frequency using bcftools. We identified cis-eQTL with QTLtools cis with the “--normal” flag within a 1 Mb window of the transcription start site. Bull age, RNA integrity number as well as the top 3 and 10 PCs, respectively, from genetic variants and RNA expression were used as fixed covariates. We performed 1000 permutations and used a false discovery rate of 5% to estimate per-gene significance thresholds, followed by a conditional pass to estimate independent eQTL signals.

Specific eQTL and nearby variants were visualised from alignment and variant call files with IGV (v2.17.4) (35).

## Results

### RNA sequencing alignment

We considered 74 mature Braunvieh bulls that had DNA-seq as well as total RNA-seq from testis, epididymis, and vas deferens tissues (4). The mean short read sequencing coverage for DNA was 13.3 ± 3.9-fold (approximately ∼240 million reads). The mean RNA-seq coverage for testis, epididymis, and vas deferens tissues was 258 ± 33, 284 ± 36, 263 ± 24 million reads respectively. After aligning the sequencing reads to the ARS-UCD1.2 bovine reference genome, an average of 99.6% of the autosomal bases were covered by at least 2 reads with DNA-seq, while for testis, epididymis, and vas deferens RNA-seq the average was 26.7%, 40.4%, and 34.8% respectively (Figure 1A). The aligned coverage for the DNA-seq reads was even across different annotated regions of the reference genome, while the RNA-seq reads were strongly enriched in protein coding regions (Figure 1B; Supplementary Figure 1). As expected for total RNA-seq, we also identified elevated coverage in regions overlapping miscellaneous micro/small/long noncoding RNA that are not typically present in mRNA-seq. We observed moderate coverage in intergenic regions, which is likely due to the incomplete annotation of the bovine genome (36), particularly of long noncoding RNAs or underrepresented tissue-specific genes.

**Figure 1.**
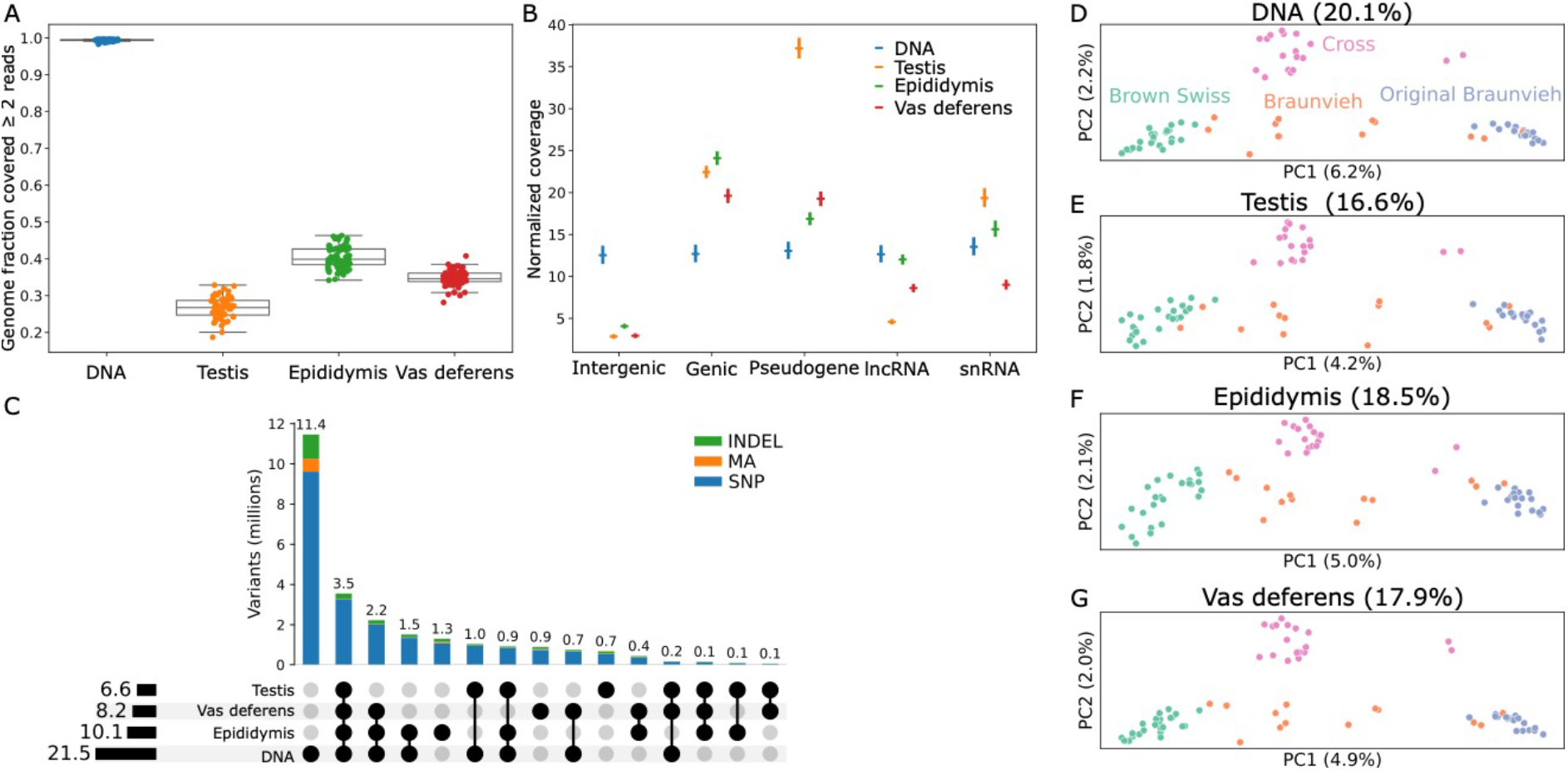
Alignment and variant calling from DNA and RNA sequencing data. (**A**) Fraction of the autosomal bases covered by at least two reads for DNA-seq and the three RNA-seq tissues. (**B**) Coverage depth normalised by the total size of the features across different annotated regions for DNA and the three RNA tissues, with respectively colours taken from (**A**). Intergenic regions have low coverage in RNA-seq while other categories are enriched, like long noncoding (lnc) and small nuclear (sn) RNA. DNA coverage is consistent across all categories. (**C**) Overlap of called variants based on exact position and REF/ALT matches, stratified by SNPs, indels, and multiallelic (MA) variants. (**D**-**G**) Principal component analyses for variants called from DNA and the three tissues. Braunvieh refers to animals of ambiguous or mixed Brown Swiss or Original Braunvieh ancestry, and cross refers to Brown Swiss or Original Braunvieh crossed with a different breed.

We then called variants using DeepVariant for each sample on the DNA and each RNA tissue type separately, using DNA- and RNA-trained models as appropriate, and then jointly genotyped the variants across all samples within each group. There were 21.5M called variants for DNA across the autosomes, and 6.6M, 8.2M, and 10.1M variants called for testis, vas deferens, and epididymis RNA respectively. The RNA samples respectively called 31%, 38%, and 47% of the number of called DNA variants (Figure 1C), which is near-proportional to the percent of the covered genome for each tissue, suggesting RNA sequencing can be used to call variants at a similar rate to DNA sequencing wherever there is coverage. Approximately 64-75% of the autosomal sequence was within 1 Kb of an RNA-called variant, compared to 95.6% for DNA-called variants (Supplementary Figure 1), implying regions of the genome remain completely inaccessible from total RNA sequencing.

The ratios of transitions to transversions (Ti:Tv) for the total RNA-seq variants ranged from 2.19 in non-exonic or noncoding exons to 3.58 within coding exons (Table 1), broadly in line with the distinct expectations for genome-wide or the more conserved coding regions (37). Most DNA-specific variants were in intergenic regions, where there was less RNA coverage. However, RNA variants within intergenic regions largely behaved as expected, although the increased Ti:Tv for epididymis and vas deferens may result from tissue-specific genes that are not yet correctly annotated. Using DNA-seq also resulted in proportionally increased indel calls, accounting for 14% of variant calls compared to ∼11% in total RNA-seq, as well as multiallelic calls (3.4% in DNA-seq versus ∼1.3% in total RNA-seq). Approximately 3.5 million variants were present in all four datasets, indicating a large portion of regions are all expressed across the three different examined tissues. We were also able to capture the same population structure using genotypes called from the three tissues as from the DNA (Figure 1D-G), demonstrating the RNA variant calls contained meaningful variation.

**Table 1.** Median number of biSNPs (biallelic SNPs) in coding exons, noncoding exons (e.g., pseudogenes, lncRNA, etc) and non-exon regions per sample with the associated Ti:Tv rate for variants called from DNA-seq or the three RNA-seq tissues.

|  | Coding exons |  | Noncoding exons |  | Not exons |  |
| --- | --- | --- | --- | --- | --- | --- |
|  | biSNPs | Ti:Tv | biSNPs | Ti:Tv | biSNPs | Ti:Tv |
| DNA | 39,657 | 3.13 | 10,113 | 2.16 | 6,652,085 | 2.20 |
| Testis | 35,458 | 3.53 | 5,669 | 2.19 | 1,427,805 | 2.20 |
| Epididymis | 34,396 | 3.53 | 6,082 | 2.21 | 2,265,030 | 2.41 |
| Vas deferens | 31,913 | 3.58 | 5,409 | 2.25 | 1,933,053 | 2.43 |

We used the variant effect predictor (VEP) software to assess potential consequences for the called variants. The RNA-seq proportionally called more variants annotated as low/moderate/high impact (Supplementary Figure 3), with the strongest enrichment (nearly 2-fold) observed in testis. On average across the tissues, between 70-75% of low/moderate/high impact variants called from DNA were present in the RNA variants, again suggesting the RNA called variants are primarily missing intergenic variants for which functional consequences are not immediately apparent.

### RNA variant calling accuracy

We examined the accuracy of RNA-seq variants, taking the DNA sequencing variants as the truth set. Although DNA-based variant calls are regarded as the gold-standard, the average depth of coverage over the 74 samples (13x) is lower than typically recommended for accurate calls (20-30x). Consequently, some false positives/negatives may be due to an imperfect truth set, particularly in heterozygous genotypes. We observed SNP precision and recall had a substantial, but expected, dependency on gene expression levels, with highly expressed genes (transcripts per million [TPM]≥10; Supplementary Figure 1) achieving 97.7% precision and 91.8% recall averaged across the three tissues, while genes with less than 0.1 TPM averaged 41.3% precision and 5.1% recall (Figure 2A). Recall in genes with less than 0.1 TPM was actually lower than that in non-exonic regions, likely due to RNA read alignments overlapping unannotated intronic or intergenic features like long noncoding RNA present in these tissues. Indel calling accuracy demonstrated a similar dependency on expression levels (Figure 2B) but with overall reduced precision and recall.

**Figure 2.**
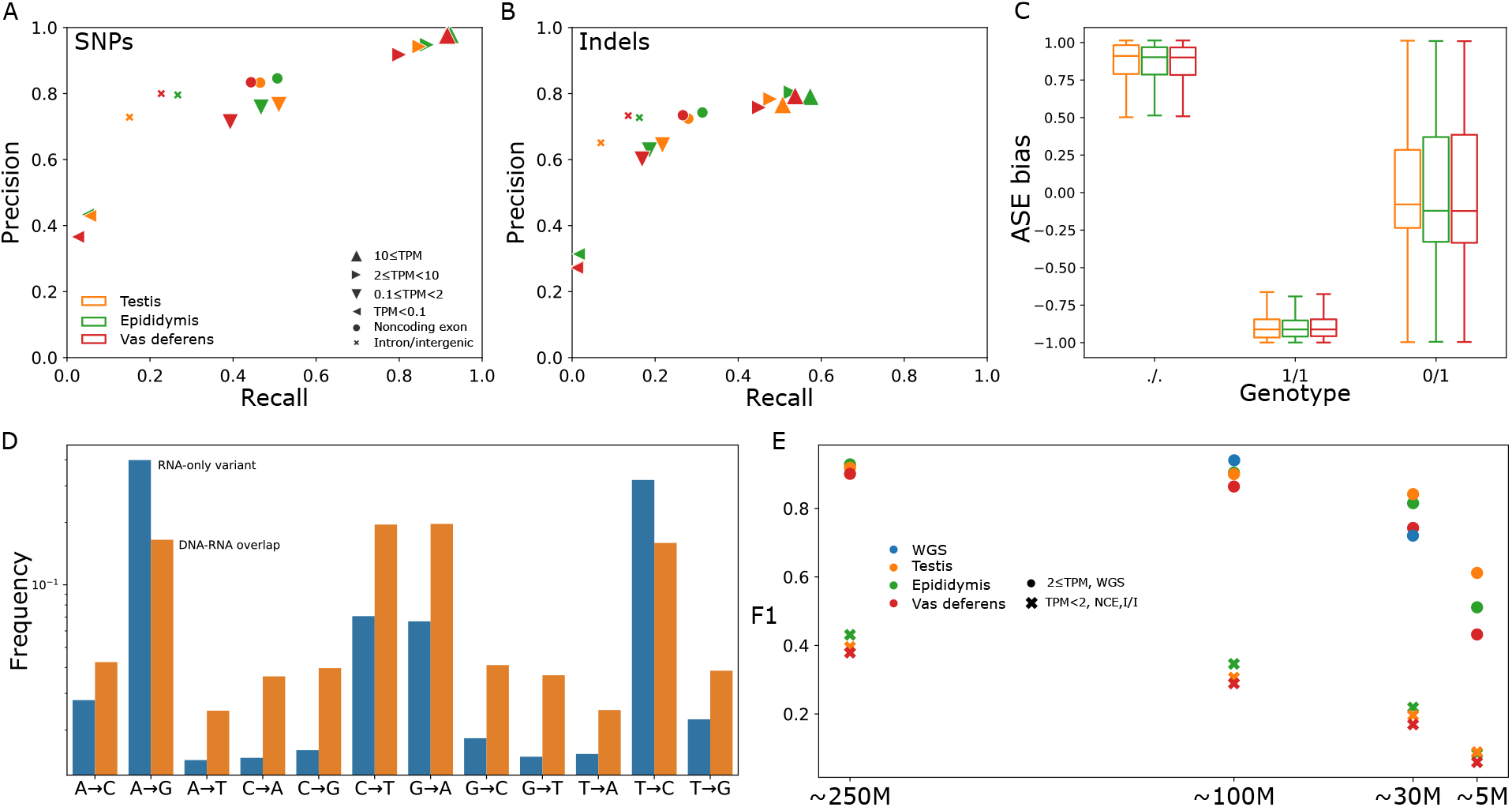
Variant precision and recall for SNPs (**A**) and indels (**B**) called from RNA-seq using DNA-seq variants as truth, stratified by non-exonic, noncoding exons, and different levels of coding exon expression. (**C**) Heterozygous variants misgenotyped as homozygous reference/missing or homozygous alternate displayed strong allelic imbalance, where positive (negative) ASE indicates the reference allele was more (less) expressed. Outliers are not plotted. (**D**) Variant calls present in all three RNA sequencing sets but not DNA were highly biased towards known patterns of RNA editing, whereas variants found in both DNA and RNA sets displayed the expected Ti:Tv behaviour. (**E**) F1 score decreases slowly as coverage is downsampled from approximately 250M reads to 100M, 30M, and 5M reads. F1 score is averaged separately for WGS or more expressed genes (TPM≥2) and less expressed genes (TPM<2), noncoding exons (NCE), or intergenic/intronic (I/I).

We also investigated the effect of allele-specific expression (ASE) on RNA-seq variant calling. Affected RNA-seq variants show a deviation from the expected 1:1 ratio of reference and alternate alleles, which negatively impacts variant calling (if the alternative allele is less expressed) and genotyping (if the reference allele is less expressed). We observe both these effects after excluding monoallelic expression, causing heterozygous DNA-seq variants to be missed or genotyped as homozygous alternate (Figure 2C). Between 56-73% of ASE-variants were still genotyped correctly, whereas mostly extreme ASE cases (>85% allelic imbalance) were erroneous.

There were 960k, 1,960k, and 1,520k variants called for testis, epididymis, and vas deferens RNA-seq, respectively, which were not called by DNA-seq. We also identified 150,011 (577,839) RNA-seq variants called uniformly across all three (two) tissues but not in the DNA-seq. Given these variants occur in different, independently sampled tissues, they potentially correspond to RNA editing or other RNA modification events that are not detectable from DNA-seq and thus appear as RNA-DNA differences (RDDs) (38), rather than erroneous variant calls. Indeed, genotyping errors attributable to ASE only explained approximately 6% of RDDs at heterozygous DNA-seq variants, and so are limited contributors to the overall observed error rate. The RDDs follow a highly biased distribution (Figure 2D), suggesting a high prevalence of A-to-I editing (A→G & T→C) and to a lesser degree C-to-U editing (G→A & C→T), two commonly reported forms of post-transcriptional RNA modifications (39). However, some of these RDDs have nearly a 100% conversion rate, suggesting this may be caused by biological mechanisms other than RNA editing or technical artefacts.

The 74 total RNA-seq samples have unusually high coverage. We downsampled the RNA-seq samples to approximately 100M, 30M, and 5M reads, roughly corresponding to typical sequencing depths suggested for splicing phenotypes, expression phenotypes, and low-pass analyses, and performed variant calling as for the full set. The fraction of autosomal sequence covered by at least two RNA reads decreased sublinearly (Supplementary Figure 2), suggesting coverage is entirely lost in lowly expressed genes but highly expressed genes are still sufficiently covered even at 5M reads. Similarly, roughly 65%, 54%, and 23% of the autosomal sequence was within 1 Kb of an RNA variant at 100M, 30M, and 5M reads (Supplementary Figure 2). The precision of called variants decreased more quickly in non-exonic or lowly expressed regions, but the precision of variants called within moderately to highly expressed exons was minimally affected down to 30M reads, and only noticeably dropped at 5M reads. Recall decreased slightly more rapidly than precision as coverage was reduced, but 30M RNA reads were still enough to capture over 70% of WGS variants in moderately to highly expressed exons. We also downsampled the DNA-seq samples to 100M and 30M reads, corresponding to genome-wide coverages of 5.3- and 1.6-fold. SNP precision and recall were slightly higher for WGS at 100M reads compared to the RNA-seq (Figure 2e). However, at 30M reads, the RNA-seq outperformed DNA-seq for both SNP precision and recall, although 1.6-fold DNA-seq is far below a typical variant calling depth and requires processing with low pass imputation approaches to achieve sufficiently accurate genotypes (40).

### eQTL mapping with DNA and RNA variants

We next investigated if the quantity and quality of variants called directly from RNA-seq is sufficient to identify expression QTL (eQTL). We conducted eQTL mapping using only the RNA-seq to genotype genomic variants and estimate molecular phenotypes and compared against a “truth set” using the conventional approach of using DNA-seq for genomic variants and RNA-seq only to estimate molecular phenotypes. We ran both permutation and conditional passes to identify independent eQTL, adjusting for covariates in both the expression matrix and variant genotypes. We assessed significance for 20,620, 21,271, and 20,097 genes expressed in testis, epididymis, and vas deferens, respectively. The RNA-only approach was able to identify 78.9%, 77.6%, and 73.6% of genes with at least one independent-acting eQTL (eGene), respectively, compared to the DNA+RNA truth approach (Figure 3A).

**Figure 3.**
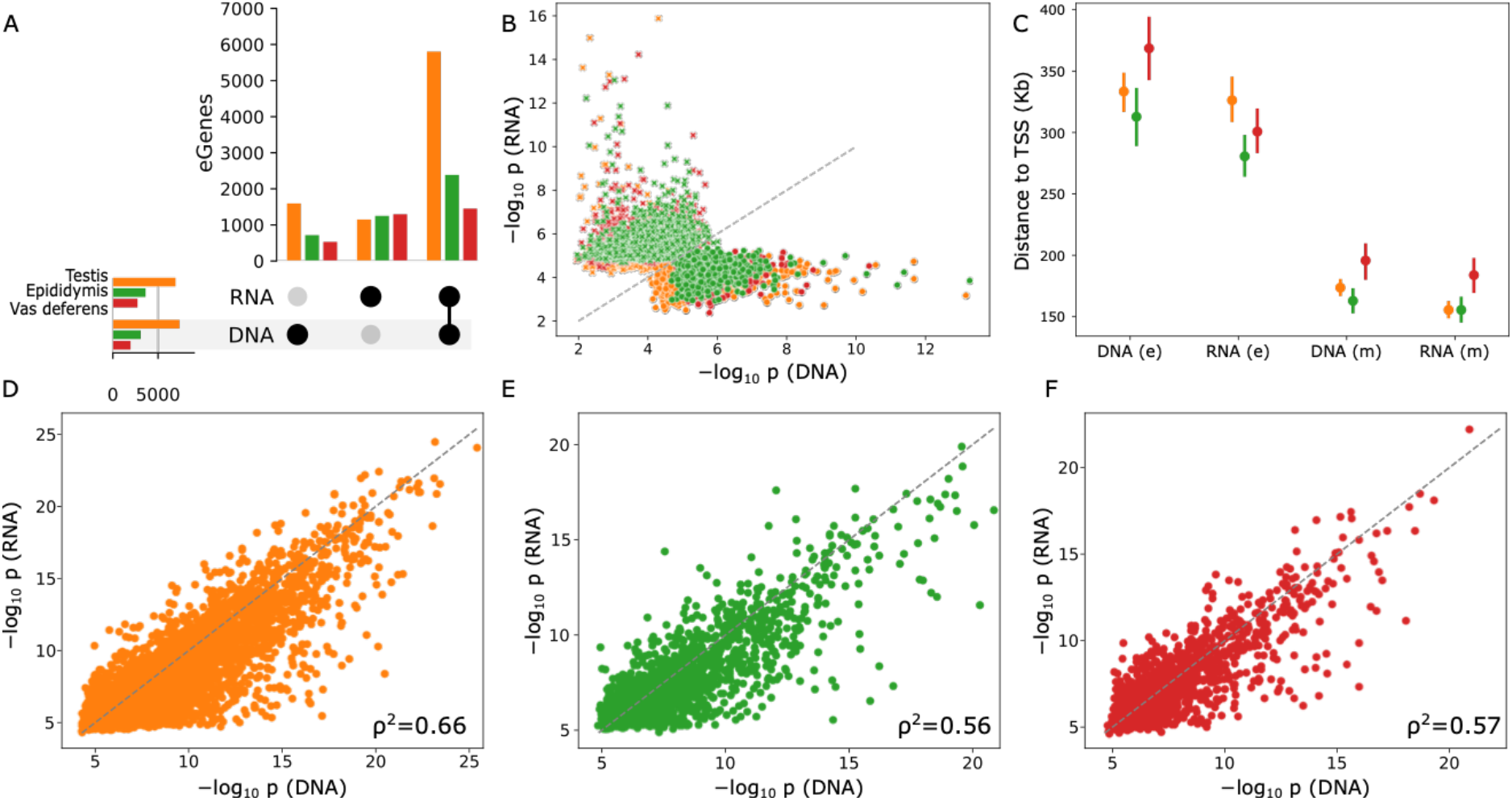
(**A**) eGenes found in both DNA and RNA variant sets or eGenes only found with RNA or DNA variants across three tissues. (**B**) The majority of eGenes found in only DNA or RNA mapping were typically close to the significance thresholds, with very few highly significant eGenes found in only one set. (**C**) eGenes found mutually (m) in both DNA and RNA sets tended to have the most significant variants closer to the TSS compared to eGenes found exclusively (e) in only one set. (**D**-**F**) P-values for the most significant variant was strongly correlated across all three tissues between the DNA and RNA variant sets for eGenes found in both.

Many of the eGenes identified exclusively in either DNA- or RNA-seq variant mapping were of lower significance and close to the discovery threshold, with the other variant set (RNA- or DNA-seq respectively) typically within an order of magnitude of the significance (Figure 3B). Only 10 and 15 unique eGenes with p-values below 1×10^-10^ were respectively found in DNA- and RNA-only association mappings. Mutual eGenes found in both DNA and RNA sets were substantially closer to the transcription start site on average (Figure 3C), as well as more significant on average compared to DNA- or RNA-only eGenes. RNA-only eGenes had substantially larger and more variable effect sizes compared to DNA-only or RNA-DNA overlapping eGenes (Supplementary Figure 4).

For RNA-DNA overlapping eGenes, we found moderate-to-strong correlation (Spearman ρ^2^ of 0.56-0.66) of the most significant p-value for each eGene when using DNA- or RNA-seq variants (Figure 3D-F). However, only approximately 9% of the RNA-DNA overlapping eGenes shared the same lead candidate variant, suggesting that while the significances were comparable, we rarely could recover the DNA-seq top eQTL using RNA-seq variants. The DNA-seq variants also had slightly more independent signal compared to using RNA-seq variants, although the effect was minor (1.13 versus 1.09 for testis, 1.07 versus 1.06 for vas deferens, and 1.03 versus 1.03 for epididymis).

We also conducted association mapping after downsampling the RNA coverage to 100M, 30M, and 5M reads, using the reduced coverage for both the RNA-seq variants and molecular phenotypes. Due to the decrease in reads used for determining gene expression, fewer genes were expressed above filtering thresholds (Supplementary Table 1), and so fewer eQTL were identified even when using the full coverage DNA-seq variants. At 100M RNA reads, there was minimal loss (1%) of QTL detection compared to using DNA-seq variants (Supplementary Figure 5), and a minor loss (5%) of detection at 30M RNA reads. Due to the substantial drop in RNA-seq variants called with 5M reads, there was a larger loss (20%) of QTL detection at this coverage relative to use DNA-seq variants.

### RNA DNA differences in eQTLs

We further examined in detail several compelling eQTL identified using only DNA- or RNA-seq variants, which contrasted to eGenes found with both sets of variants (e.g., *ENSBTAG00000000261;* Figure 4A, B) in terms of RNA-seq variant density and imputation accuracy. *ENSBTAG00000000597* was a strongly associated eGene in epididymis when using DNA-seq variants (p=5.3×10^-14^), but not significant with RNA-seq variants (p=1.4×10^-4^). The same top SNP variant was called in both DNA and RNA variant sets (Figure 4C, D), but was poorly genotyped in epididymis RNA (allele frequency of 0.26 in DNA-seq and 0.07 in RNA-seq) resulting from a low *ENSBTAG00000000597* transcript abundance (average TPM 0.23). Only several of the epididymis samples even had RNA-seq coverage around this region, leading to extremely imbalanced genotypes for all variants in a 1 Kb window, with a significant deviation from Hardy-Weinberg proportions (p=9.2×10^-7^) while the DNA-seq variants followed Hardy-Weinberg proportions (p=0.86). Consequently, no significant association between RNA-called variants and *ENSBTAG00000000597* expression was found. Similarly, an eQTL for *ENSBTAG00000033056* was missed in testis, within only 5 low quality variants within a 5 Kb window of the lead DNA SNP. In general, almost all DNA-only eQTL were due to the lack of well genotyped RNA variants near the lead DNA SNP. We did not observe any DNA-only QTL where the missing RNA variants could be explained by ASE.

**Figure 4.**
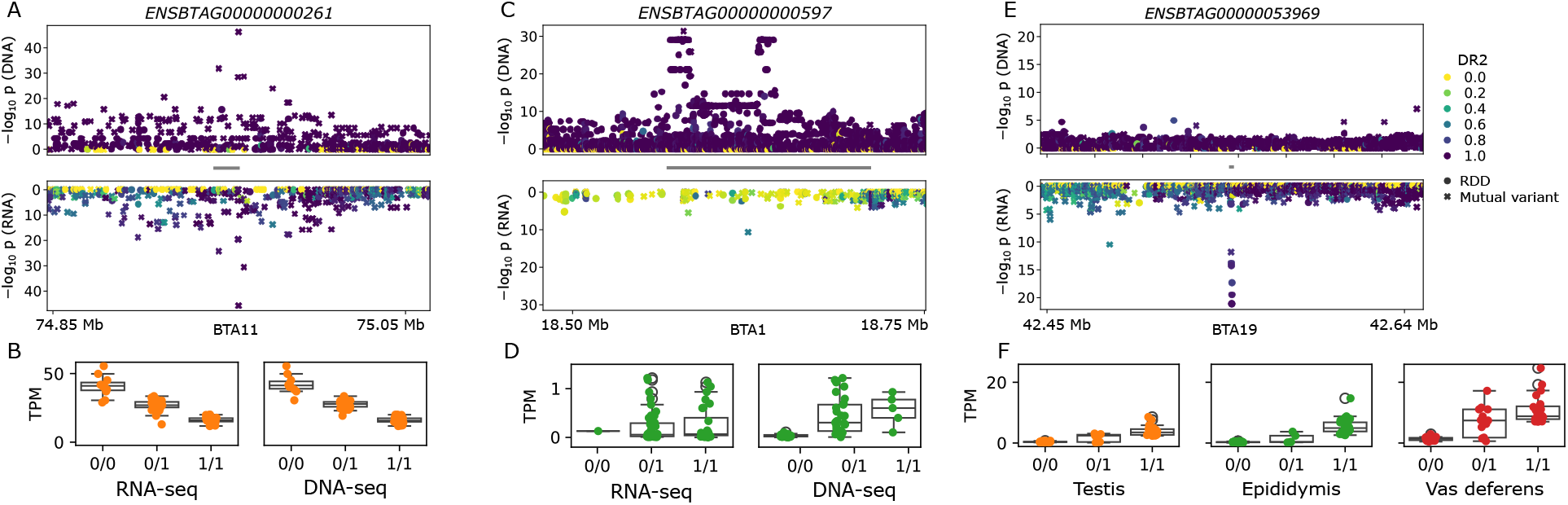
(**A, C, E**) Zoom plots for an eGene identified with both variant sets, DNA-seq variants only, and RNA-seq variants only respectively. The grey bar between the DNA and RNA associations represents the gene, while the marker colour represents imputation accuracy (DR2). The marker style indicates if the variant is present in both DNA-seq and RNA-seq variants or if it is an RDD. (**B, D, F**) TPM plots for their respective three genes. The same lead variant is used for as the genotype in **B** and **D** for testis and epididymis respectively, while the lead variant for **C** is an RDD and can only be examined for RNA-seq but is present in all three tissues.

Unexpectedly, some eGenes are only identified when mapping RNA-seq variants and not with DNA-seq variants. For example, *ENSBTAG00000020116* was significant in epididymis tissue, but primarily because the conditional significance threshold was moderately lower for RNA-seq variants. Fewer RNA-seq variants within the cis-window led to a significance threshold of 3.5×10^-6^ (versus a DNA threshold of 7.9×10^-7^), and the top RNA variant had p=2.2×10^-6^ (versus DNA top variant p=5.0×10^-6^). These marginal examples could be removed by setting a uniform stricter significance threshold, especially in the case of sparse variants.

Out of the 15 highly significant (p<1×10^-10^) RNA-seq only eGenes, only nine are annotated as protein coding, while the other six are e.g., pseudogenes or lncRNA (Supplementary Table 2). Genome-wide, protein coding genes make up 80% of the annotation, compared to only 60% of these RNA-only eGenes. Most of these highly significant RNA-seq only eGenes appear in all three examined tissues, suggesting this is not a tissue-specific observation but potentially something affecting RNA analyses more generally. Almost all of these genes have multiple paralogues (Supplementary Table 2), which can lead to low-quality or ambiguous RNA alignments and thus degraded variant calling. However, we find, for example in *ENSBTAG00000053969* (Figure 4E, F) and *ENSBTAG00000027962* (Supplementary Figure 6), that RNA-seq coverage can be largely missing or highly expressed in a portion of the annotated exon region (Supplementary Figure 7). The top associated variants appear within these differentially covered regions, and some homozygous reference samples have sufficient coverage to be distinguished from a missing genotype. The lack of variants in the DNA-seq and the distinct RNA coverage dropout suggest these eQTL cannot simply be explained by paralogue mismapping for the RNA reads, although it is not clear if there is an alternative artefactual explanation or a mechanism beyond the genome (e.g., RNA editing/modification, epigenome, etc.)

## Discussion

RNA sequencing is critical to examine mechanisms underpinning variation in gene expression or splicing but its utility for variant calling had not been characterised extensively. We find deep total RNA sequencing with ∼250M reads covers one third of the genome, leaving many (primarily intergenic) regions inaccessible. From 74 cattle transcriptomes, we call 7-10M variants per tissue, approximately only 40% of that from matched DNA sequencing, but still two orders of magnitude more than previously reported for cattle RNA variant calling from primarily mRNA (9). Particularly in coding regions that are highly expressed (TPM≥10), we recover over 92% of DNA-seq variants with precision of approximately 98%. Precision and recall are reduced at more typical RNA coverage levels, with 76% precision and 26% recall genome-wide at 30M reads. Testis (41), and likely epididymis and vas deferens as well (4), express substantially more genes at detectable levels compared to other tissues, meaning that our recall values likely represent an upper bound and might be lower for most other tissues. RNA-specific effects, like allele-specific expression or RNA editing, are detrimental to variant calling accuracy but only affect a limited number of sites.

Despite total RNA-seq variant calling only capturing approximately 40% of variant sites compared to DNA-seq variant calling, it identifies roughly 75% of eGenes, and so is nearly 2-fold enriched for eQTL. This trend holds when downsampling to 30M reads before sharply dropping at 5M reads. Interestingly, when downsampling to 30M reads, we find only 10-15% fewer expressed genes but roughly 50% fewer significant eGenes (Supplementary Table 1), suggesting that deeper coverage is required for comprehensively mapping eQTL (4). The majority of eGenes identified by DNA-seq but missed by RNA-seq variants are due to eQTL being extremely distant to the TSS (>300 Kb) or, to a lesser degree, located within lowly expressed regions leading to poor RNA variant genotyping accuracy. On the other hand, highly significant eGenes unique to RNA-seq variants are mostly associated with RDDs (12 out of 15) with few variants in linkage disequilibrium, which would likely fail manual curation. However, the leading RNA-only eQTL variants have extremely high variant qualities and imputation scores, and so cannot be easily filtered *a priori*. Furthermore, the low agreement we observed for the top associated variants between DNA-seq and RNA-seq variants would weaken downstream analyses like colocalization of putative causal variants (4) if depending only on RNA-seq variants.

Livestock GTEx consortia rely on RNA-seq for variant calling (e.g., cattle (8), pig (12), and chicken (11)) to enable molecular QTL mapping as most RNA-sequenced samples don’t have matched DNA-based genotypes or sequences. This is different to the equivalent human GTEx (13) which uses transcriptomes that have matched DNA whole-genome sequencing. We have comprehensively shown that RNA-seq variant calling accuracy is highly dependent on gene expression levels (∼98% precision for TPM≥10 versus ∼75% precision for 2>TPM≥0.1) and hundreds of thousands of RDDs exist in each tissue examined. While these livestock GTExs impute RNA-seq variants into large reference panels, likely avoiding the false positive RNA-seq variant eGenes, our results demonstrate that caution is needed when using RNA-seq variants as a replacement for DNA-seq or genotyping array variants.

Considerable uncertainty remains over the origins of RDDs, and whether they are technical artefacts or biological modifications (42–44). Over 150k variants are called in all three RNA-seq tissues but not in DNA-seq, many of which had high allele frequencies and allele depth, demonstrating that RDDs are pervasive regardless of their true origin, and cannot be simply addressed by conservative filtering. Analogous to improvements in alignment uniqueness for long read over short read DNA (45), aligning to genes with highly similar paralogues likely will likely benefit from long read RNA approaches and disentangle potential causes like paralogue or pseudogene alignments (46) for some RDDs.

## Data availability

There are no new data associated with this article. All scripts and pipelines used in these analyses are available at https://github.com/AnimalGenomicsETH/RNA_variant_calling.

## Author Contributions

ASL: Conceptualization, Investigation, Visualisation, Writing—original draft. XMM: Writing—Validation, review & editing. HP: Supervision, Writing—review & editing

