## Supplementary material for "RNA sequencing variants are enriched for eQTL in cattle tissues"

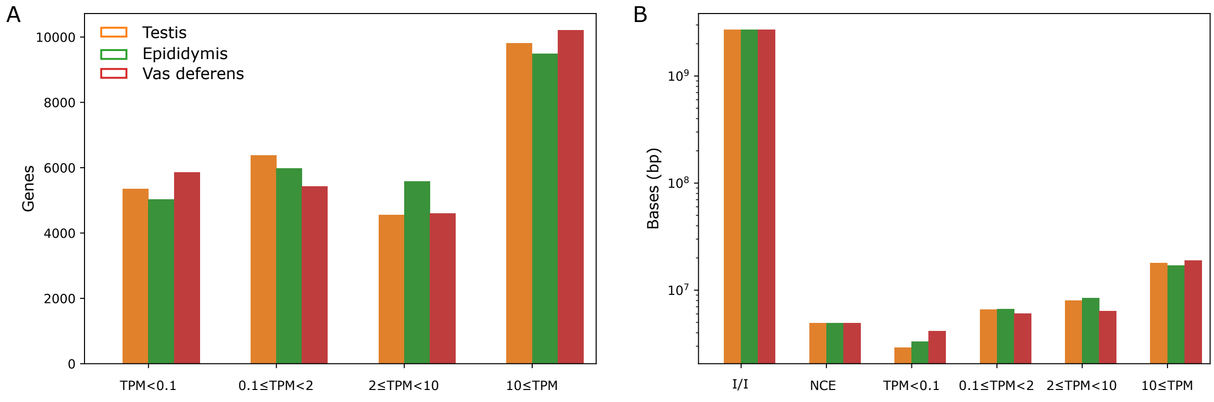


*Supplementary Figure 1. (**A**) Number of genes expressed at different TPM thresholds. (**B**) Total number of bases classified into different regions, for intergenic/intronic (I/I), noncoding exons (NCE), and the different TPM thresholds for coding exons.*

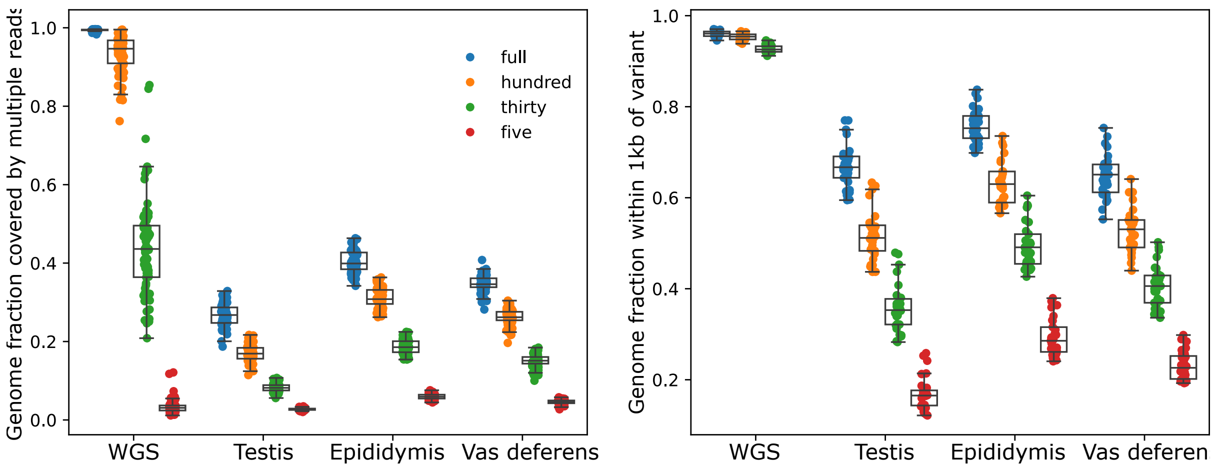


*Supplementary Figure 2. (**A**) Fraction of autosomes covered by at least two reads at different levels of subsampling. (**B**) Fraction of autosomal sequence with at least one variant within 1 Kb at different levels of subsampling.*

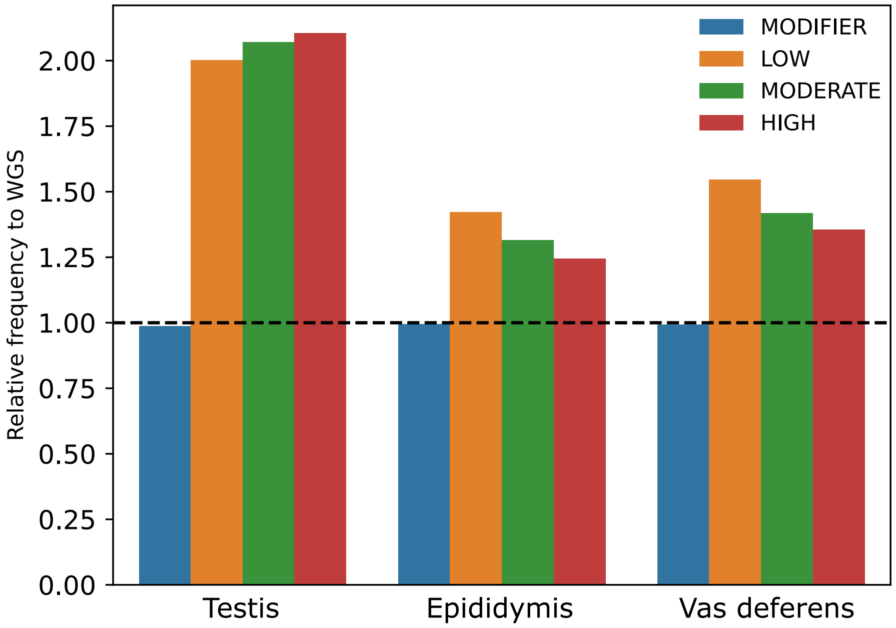


*Supplementary Figure 3. Relative frequency of Variant Effect Predictor "IMPACT" feature normalised per impact feature relative to WGS.*

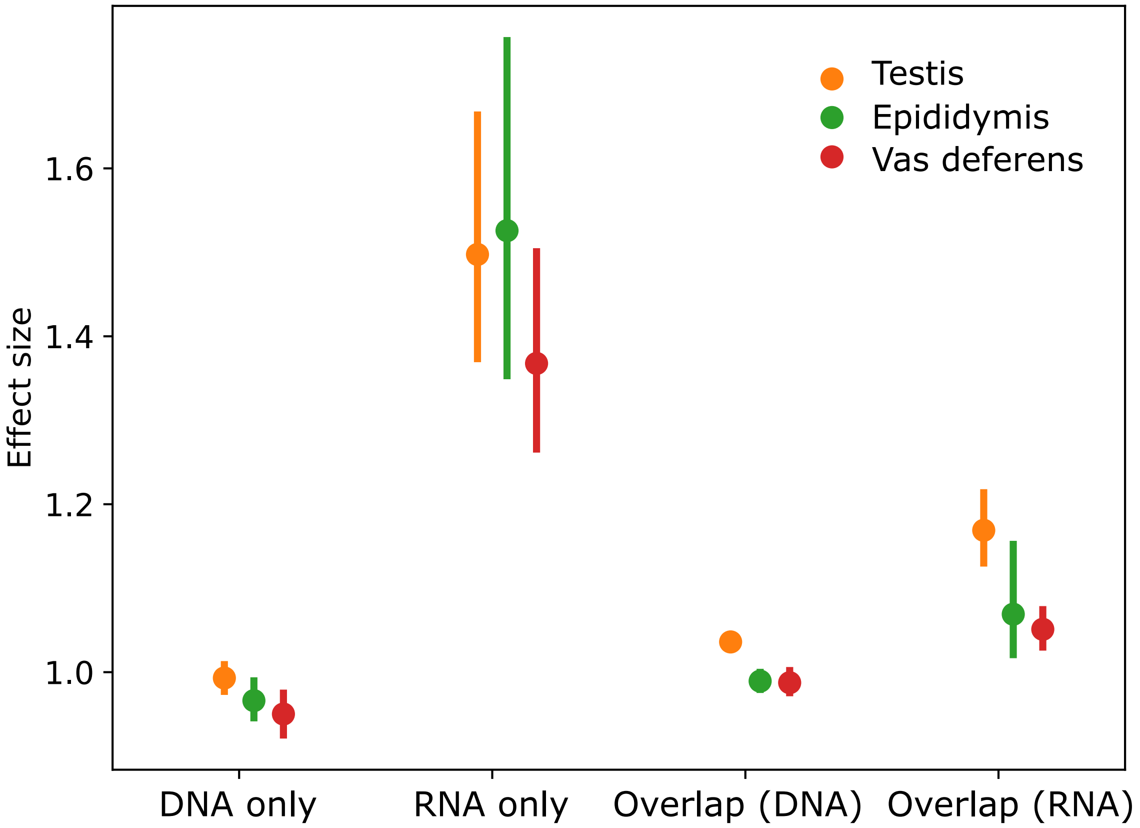


*Supplementary Figure 4. Absolute values for effect size for eQTL in three tissues for eQTL found uniquely with DNA variants or RNA variants, as well as common eQTL.*

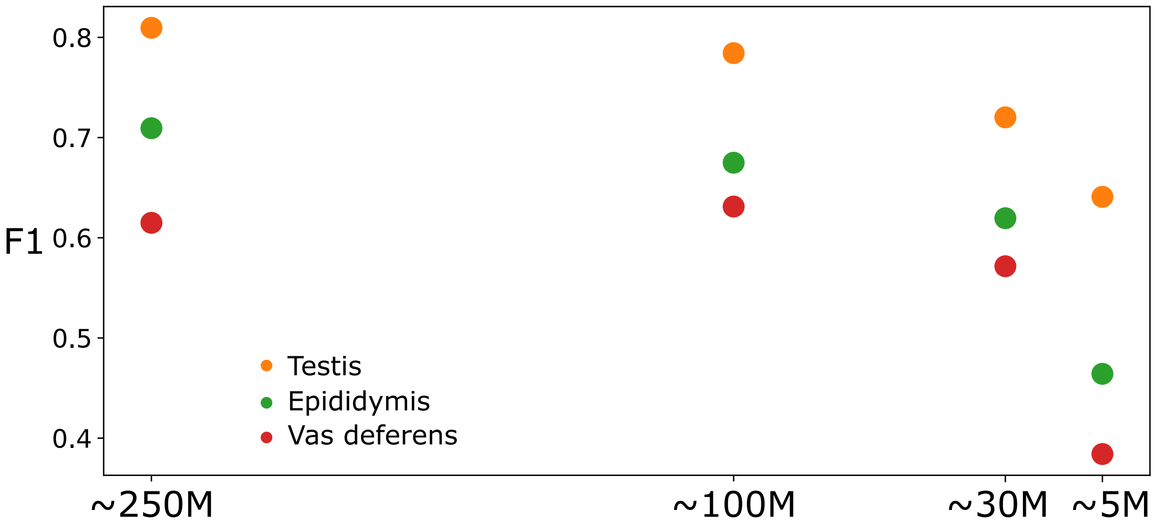


*Supplementary Figure 5. F1 score taking DNA-variant eGenes as truth. Molecular phenotypes are also calculated from the subsampled RNA coverage, and so the number of eGenes in the truth set also shrinks at lower coverage.*

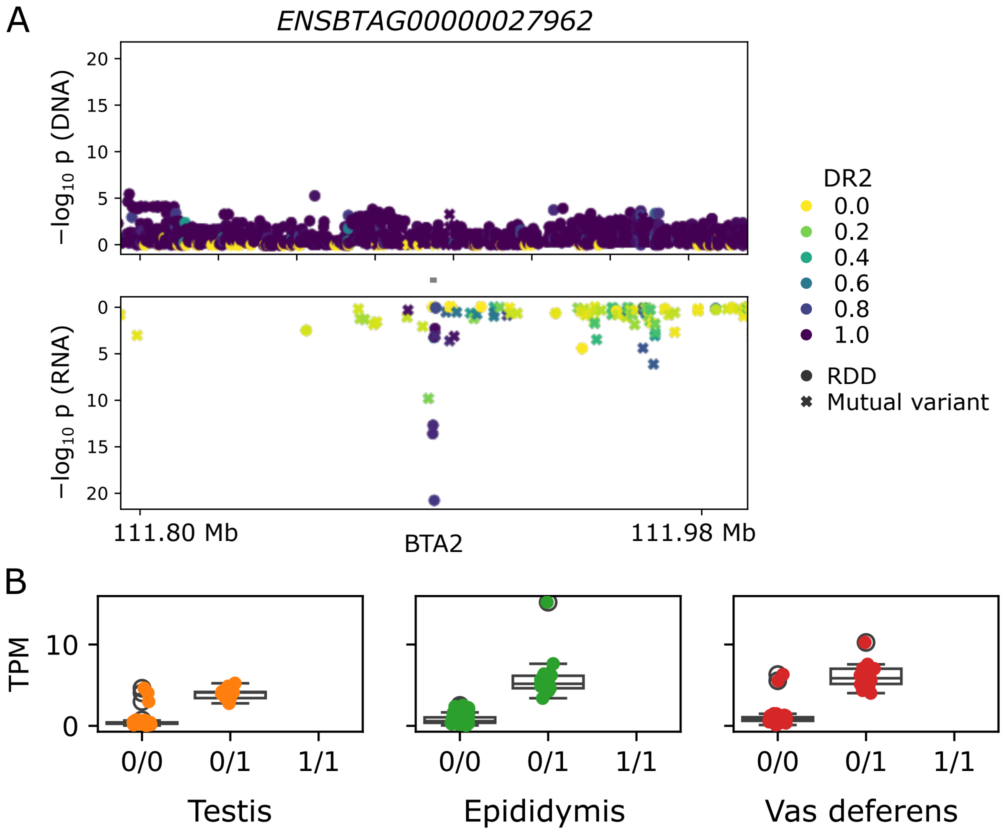


*Supplementary Figure 6. (**A**) Zoom plots for an eGene associated only with an RNA-seq variant. The grey bar between the DNA and RNA associations represents the gene, while the marker colour represents imputation accuracy (DR2). The marker style indicates if the variant is present in both DNA-seq and RNA-seq variants or if it is an RDD. (**B**) TPM plot for the respective gene. The lead variant is an RDD and can only be examined for RNA-seq but is present in all three tissues.*

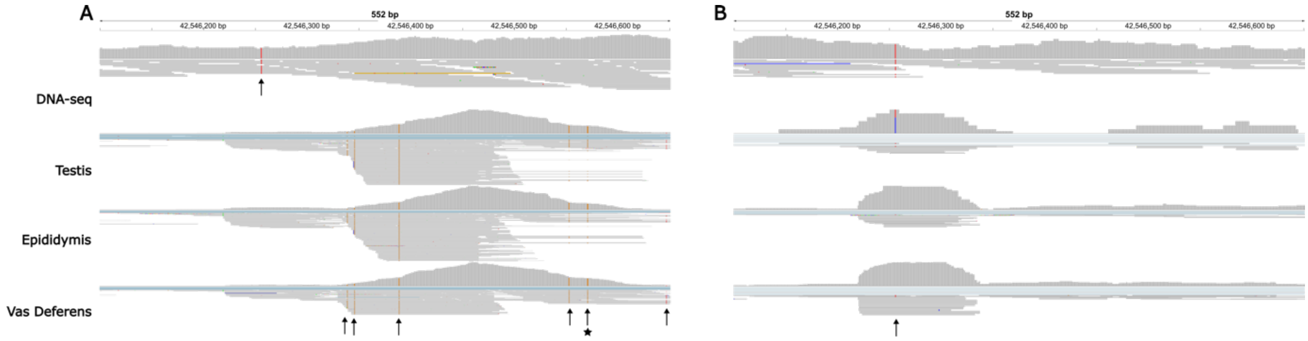


*Supplementary Figure 7. IGV alignments of DNA-seq and RNA-seq for the three tissues surrounding the lead variant for* ENSBTAG00000053969 *in a sample with homozygous alternate genotypes (**A**) and homozygous reference genotypes (**B**) over the range 19:42546100-42546650. RDDs are marked by arrows, and the lead eQTL variant is additionally marked with a star. The RNA-seq coverage scales are significantly different, ranging up to 800 aligned reads in (**A**) and up to 70 aligned reads in (**B**).*

*Supplementary Table 1. Number of expressed genes passing filtering thresholds for the three RNA tissues at full coverage (~250 M reads) as well as the three downsampled coverages. The number of significant eGenes (using DNA-seq variants) is given in parentheses.*

| | Full coverage | 100 M reads | 30 M reads | 5 M reads |
|---|---|---|---|---|
| Testis | 20,960 (7,377) | 20,169 (6,229) | 18,563 (4,220) | 15,867 (1,264) |
| Epididymis head | 21,271 (3,084) | 20,260 (2,443) | 18,423 (1,227) | 15,272 (234) |
| Vas deferens | 20,097 (1,962) | 19,032 (1,473) | 17,065 (965) | 13,936 (239) |

*Supplementary Table 2. eGenes uniquely detected with RNA-seq and not with DNA-seq as the source of genomic variants. Many of the eGenes have substantial number of paralogues or pseudogenes, leading to higher likelihood of mis/multi-mapping reads. Tissues indicated with * mean the eGene was uniquely found using RNA-seq variants, but the significance was not below $1{\times}10^{-10}$. Top associated variants are indicated as RNA-DNA differences (RDD) or not.*

| Gene | Gene type | paralogues | Tissues | RDD |
|---|---|---|---|---|
| *ENSBTAG00000001858* | Protein coding | 5 | T, E, V | Y |
| *ENSBTAG00000007816* | Processed pseudogene | 3 | E, V | Y |
| *ENSBTAG00000009110* | Pseudogene | 1 | T*, E*, V | Y |
| *ENSBTAG00000012798 (KCNH8)* | Protein coding | 17 | E*, V | N |
| *ENSBTAG00000020620 (RGS2)* | Protein coding | 25 | T*, E*, V | Y |
| *ENSBTAG00000021565 (PRSS2)* | Protein coding | 3 | T | N |
| *ENSBTAG00000027962* | Protein coding | 1 | T, E, V | Y |
| *ENSBTAG00000038064* | Protein coding | 5 | T, E | N |
| *ENSBTAG00000040518* | Protein coding | 22 | T*, E, V* | Y |
| *ENSBTAG00000042358* | snoRNA | 16 | T*, E, V* | Y |
| *ENSBTAG00000043349* | snoRNA | 16 | T, E, V* | Y |
| *ENSBTAG00000050214* | Processed pseudogene | 6 | T, E*, V* | Y |
| *ENSBTAG00000053419* | Protein coding | 2 | T*, E, V* | Y |
| *ENSBTAG00000053570* | Processed pseudogene | 22 | T*, E, V | Y |
| *ENSBTAG00000053969* | Protein coding | 21 | T, E, V | Y |

*Supplementary Table 3. Publicly available DNA-seq and matched RNA-seq in three tissues for 74 Braunvieh cattle samples, with approximately 10-fold coverage for each DNA-seq sample and approximately 250M paired-end reads for the RNA-seq samples.*

| Sample | DNA | Testis | Epididymis | Vas deferens |
|---|---|---|---|---|
| BV_1 | SAMEA9539940 | SAMEA9540537 | SAMEA113601208 | SAMEA113601309 |
| BV_2 | SAMEA9539941 | SAMEA9540538 | SAMEA113601233 | SAMEA113601330 |
| BV_3 | SAMEA111328928 | SAMEA111328928 | SAMEA113601279 | SAMEA113601366 |
| BV_4 | SAMEA111328929 | SAMEA111328929 | SAMEA113601280 | SAMEA113601367 |
| BV_5 | SAMEA111328925 | SAMEA111328925 | SAMEA113601276 | SAMEA113601361 |
| BV_6 | SAMEA9539945 | SAMEA9540542 | SAMEA113601252 | SAMEA113601346 |
| BV_7 | SAMEA9539946 | SAMEA9540543 | SAMEA113601232 | SAMEA113601329 |
| BV_8 | SAMEA9539947 | SAMEA9540544 | SAMEA113601212 | SAMEA113601312 |
| BV_9 | SAMEA9539948 | SAMEA9540545 | SAMEA113601213 | SAMEA113601314 |
| BV_10 | SAMEA9539949 | SAMEA9540546 | SAMEA113601215 | SAMEA113601316 |
| BV_11 | SAMEA111328926 | SAMEA111328926 | SAMEA113601277 | SAMEA113601362 |
| BV_12 | SAMEA111328917 | SAMEA111328917 | SAMEA113601266 | SAMEA113601355 |
| BV_13 | SAMEA113578975 | SAMEA113578975 | SAMEA113601290 | SAMEA113601375 |
| BV_14 | SAMEA9539950 | SAMEA9540547 | SAMEA113601250 | SAMEA113601345 |
| BV_15 | SAMEA9539953 | SAMEA9540550 | SAMEA113601244 | SAMEA113601340 |
| BV_16 | SAMEA9539954 | SAMEA9540551 | SAMEA113601220 | SAMEA113601317 |
| BV_17 | SAMEA9539955 | SAMEA9540552 | SAMEA113601214 | SAMEA113601315 |
| BV_18 | SAMEA9539957 | SAMEA9540554 | SAMEA113601248 | SAMEA113601343 |
| BV_19 | SAMEA9539961 | SAMEA9540558 | SAMEA113601205 | SAMEA113601306 |
| BV_20 | SAMEA9539964 | SAMEA9540561 | SAMEA113601204 | SAMEA113601305 |
| BV_21 | SAMEA9539966 | SAMEA9540563 | SAMEA113601195 | SAMEA113601298 |
| BV_22 | SAMEA9539967 | SAMEA9540564 | SAMEA113601241 | SAMEA113601337 |
| BV_23 | SAMEA9539968 | SAMEA9540565 | SAMEA113601201 | SAMEA113601303 |
| BV_24 | SAMEA9539969 | SAMEA9540566 | SAMEA113601239 | SAMEA113601334 |
| BV_25 | SAMEA111328924 | SAMEA111328924 | SAMEA113601275 | SAMEA113601360 |
| BV_26 | SAMEA111328941 | SAMEA111328941 | SAMEA113601294 | SAMEA113601380 |
| BV_27 | SAMEA113578979 | SAMEA113578979 | SAMEA113601284 | SAMEA113601372 |
| BV_28 | SAMEA113578980 | SAMEA113578980 | SAMEA113601291 | SAMEA113601377 |
| BV_29 | SAMEA9539970 | SAMEA9540567 | SAMEA113601230 | SAMEA113601327 |
| BV_30 | SAMEA9539971 | SAMEA9540568 | SAMEA113601199 | SAMEA113601301 |
| BV_31 | SAMEA111328930 | SAMEA111328930 | SAMEA113601282 | SAMEA113601369 |
| BV_32 | SAMEA113578981 | SAMEA113578981 | SAMEA113601278 | SAMEA113601365 |
| BV_33 | SAMEA9539972 | SAMEA9540569 | SAMEA113601243 | SAMEA113601339 |
| BV_34 | SAMEA9539974 | SAMEA9540571 | SAMEA113601240 | SAMEA113601336 |
| BV_35 | SAMEA9539975 | SAMEA9540572 | SAMEA113601198 | SAMEA113601300 |
| BV_36 | SAMEA9539976 | SAMEA9540573 | SAMEA113601200 | SAMEA113601302 |
| BV_37 | SAMEA9539977 | SAMEA9540574 | SAMEA113601229 | SAMEA113601326 |
| BV_38 | SAMEA9539978 | SAMEA9540575 | SAMEA113601223 | SAMEA113601320 |
| BV_39 | SAMEA9539979 | SAMEA9540576 | SAMEA113601227 | SAMEA113601324 |

| | | | | |
|---|---|---|---|---|
| BV_40 | SAMEA113578967 | SAMEA113578967 | SAMEA113601272 | SAMEA113601358 |
| BV_41 | SAMEA111328932 | SAMEA111328932 | SAMEA113601285 | SAMEA113601373 |
| BV_42 | SAMEA111328931 | SAMEA111328931 | SAMEA113601283 | SAMEA113601370 |
| BV_43 | SAMEA9539980 | SAMEA9540577 | SAMEA113601231 | SAMEA113601328 |
| BV_44 | SAMEA9539981 | SAMEA9540578 | SAMEA113601207 | SAMEA113601308 |
| BV_45 | SAMEA9539982 | SAMEA9540579 | SAMEA113601211 | SAMEA113601313 |
| BV_46 | SAMEA111328916 | SAMEA111328916 | SAMEA113601265 | SAMEA113601354 |
| BV_47 | SAMEA111328913 | SAMEA111328913 | SAMEA113601263 | SAMEA113601352 |
| BV_48 | SAMEA9539984 | SAMEA9540581 | SAMEA113601202 | SAMEA113601304 |
| BV_49 | SAMEA9539986 | SAMEA9540583 | SAMEA113601210 | SAMEA113601311 |
| BV_50 | SAMEA113578969 | SAMEA113578969 | SAMEA113601281 | SAMEA113601368 |
| BV_51 | SAMEA9539989 | SAMEA9540586 | SAMEA113601242 | SAMEA113601338 |
| BV_52 | SAMEA9539990 | SAMEA9540587 | SAMEA113601197 | SAMEA113601299 |
| BV_53 | SAMEA9539991 | SAMEA9540588 | SAMEA113601221 | SAMEA113601318 |
| BV_54 | SAMEA9539992 | SAMEA9540589 | SAMEA113601225 | SAMEA113601322 |
| BV_55 | SAMEA9539993 | SAMEA9540590 | SAMEA113601226 | SAMEA113601323 |
| BV_56 | SAMEA111328912 | SAMEA111328912 | SAMEA113601262 | SAMEA113601351 |
| BV_57 | SAMEA9539999 | SAMEA9540596 | SAMEA113601209 | SAMEA113601310 |
| BV_58 | SAMEA9540000 | SAMEA9540597 | SAMEA113601228 | SAMEA113601325 |
| BV_59 | SAMEA9540005 | SAMEA9540602 | SAMEA113601249 | SAMEA113601344 |
| BV_60 | SAMEA9540002 | SAMEA9540599 | SAMEA113601247 | SAMEA113601342 |
| BV_61 | SAMEA113578970 | SAMEA113578970 | SAMEA113601289 | SAMEA113601374 |
| BV_62 | SAMEA9540001 | SAMEA9540598 | SAMEA113601224 | SAMEA113601321 |
| BV_63 | SAMEA111328940 | SAMEA111328940 | SAMEA113601293 | SAMEA113601379 |
| BV_64 | SAMEA9540003 | SAMEA9540600 | SAMEA113601245 | SAMEA113601341 |
| BV_65 | SAMEA9540007 | SAMEA9540604 | SAMEA113601255 | SAMEA113601348 |
| BV_66 | SAMEA9540008 | SAMEA9540605 | SAMEA113601253 | SAMEA113601347 |
| BV_67 | SAMEA9540009 | SAMEA9540606 | SAMEA113601238 | SAMEA113601333 |
| BV_68 | SAMEA113578971 | SAMEA113578971 | SAMEA113601273 | SAMEA113601359 |
| BV_69 | SAMEA9540011 | SAMEA9540608 | SAMEA113601237 | SAMEA113601332 |
| BV_70 | SAMEA111328920 | SAMEA111328920 | SAMEA113601269 | SAMEA113601356 |
| BV_71 | SAMEA111328922 | SAMEA111328922 | SAMEA113601271 | SAMEA113601357 |
| BV_72 | SAMEA9540013 | SAMEA9540610 | SAMEA113601257 | SAMEA113601349 |
| BV_73 | SAMEA9540015 | SAMEA9540612 | SAMEA113601235 | SAMEA113601331 |
| BV_74 | SAMEA111328942 | SAMEA111328942 | SAMEA113601295 | SAMEA113601381 |
